## Supplementary material for "Balancing safety and efficiency in human decision making": Tracked changes for revised manuscript

Fontanesi et al., 2019, Fengler et al., 2022].

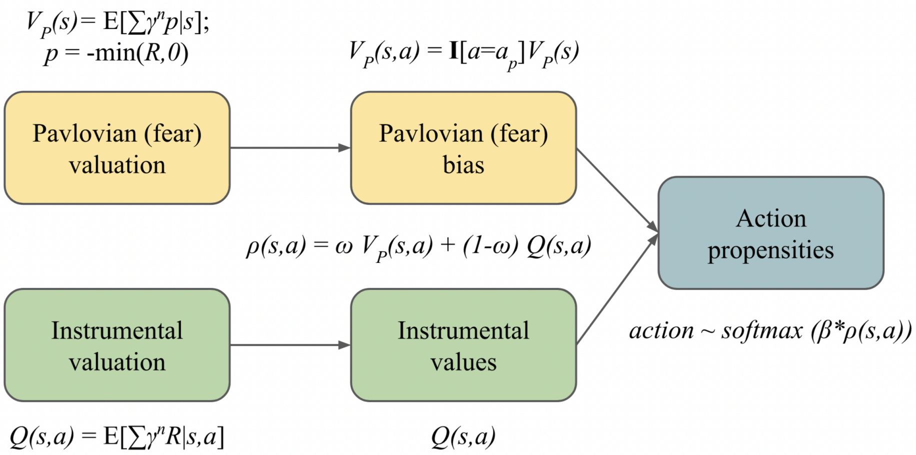


**Figure 1.** Pavlovian and Instrumental valuations are combined to arrive at action propensities used for (softmax) action selection. The Pavlovian bias influences protective behaviours through safer (Boltzmann) exploration and the arbitration between the Pavlovian and Instrumental systems is performed using the parameter $\omega$. Here $R$ denotes the feedback signal which can take both positive values (in the case of rewards) and negative values (in the case of punishments). Please see Methods for technical details; notations for the illustration follow Dorfman and Gershman [2019].

However, the flexible omega policy (with $\alpha_\Omega = 0.6$ and $\kappa = 6.5$) achieves safety almost comparable to $\omega = 0.5$ (which is the safest fixed $\omega$ policy amongst $\omega = 0.1, 0.5, 0.9$ at a much higher efficiency than $\omega = 0.5, 0.9$, thus improving the safety-efficiency trade-offs (Fig. 2F). In this way, PAL model encourages cautious exploration early on when uncertainty is higher and reduces the Pavlovian biases as the uncertainty is resolved (Fig. 2F). The flexible $\omega$ value at convergence depends on the environment statistics: transition probabilities and reward/punishment magnitudes. We utilise a simple linear scaling of associability clipped at 1 to arrive at arbitrator $\omega$ (equation 15) instead of another alternative such as sigmoid to avoid additional unnecessary meta parameters (i.e. bias shift) to be tuned. In this environment, the value at convergence is $\omega = 0.42$, due to some irreducible uncertainty in state transitions (10% chance of incorrect transition). The differences in learned instrumental value functions between PAL and a purely instrumental agent are visible in (Fig. 2C) showing how the Pavlovian bias sculpts instrumental value acquisition.

In the supplementary methods, we provide additional simulations that show the robustness of these results with respect to metaparameters $\alpha_\Omega$ and $\kappa$ (Appendix A.1), environments in which the reward locations vary (Appendix A.2), and other grid-world environments (Appendix A.3).

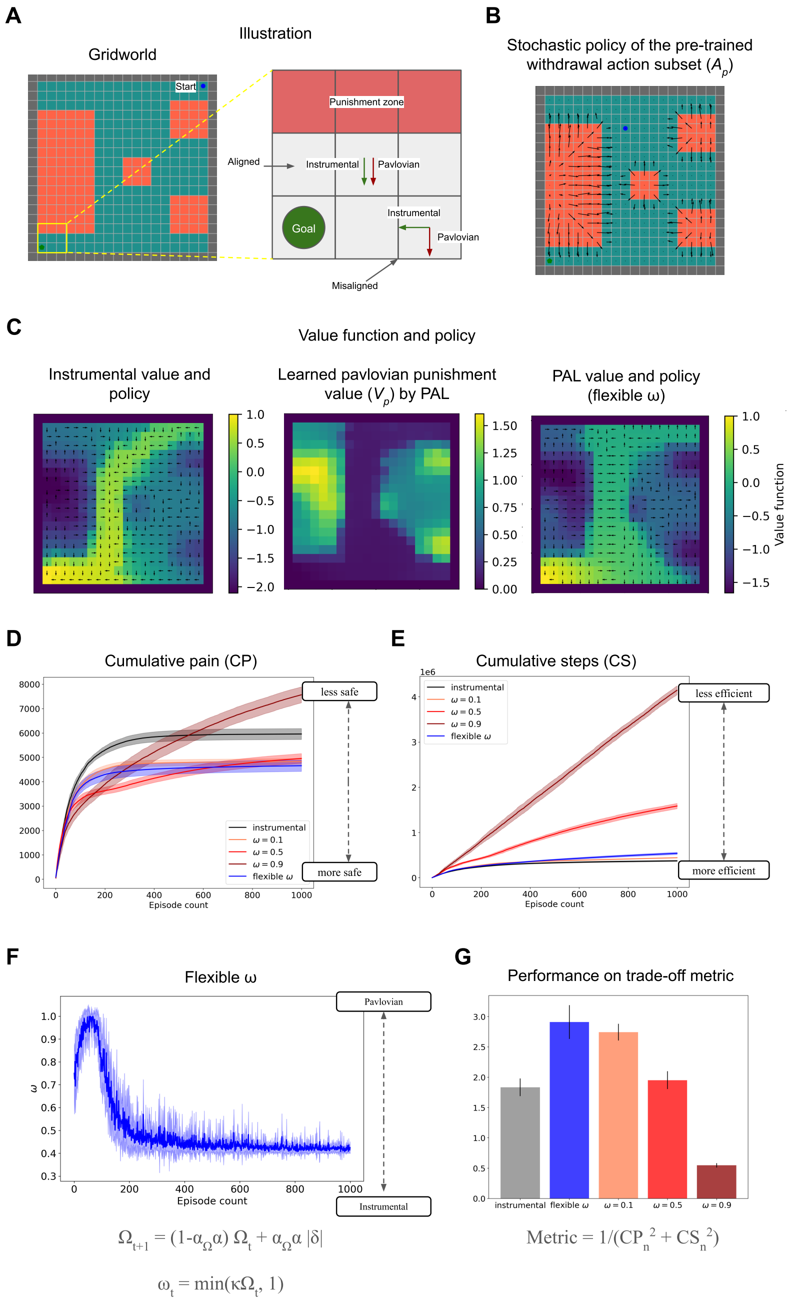


**Figure 2.** (A) Grid world environment with starting state in the top-right corner and rewarding goal state ($R = +1$) in the bottom-left corner and the red states are painful ($R = -0.1$). The grid world layout follows Gehring and Precup [2013]. Inset provides a didactic example of misalignment between Pavlovian bias and Instrumental action. (B) Stochastic policy of pre-trained withdrawal action subset $A_p$, which is biased with Pavlovian punishment values in the PAL agent. (C) The learned instrumental values and Pavlovian fear bias $V_p$(heatmap) and policy (arrows) are learned by the instrumental and flexible $\omega$ agent by the end of the learning duration. The value functions plotted are computed in an on-policy manner. (D) Cumulative pain accrued by fixed and flexible $\omega$ agents whilst learning over 1000 episodes as a measure of safety averaged over 10 runs. (E) Cumulative steps required to reach the fixed goal by fixed and flexible $\omega$ agents whilst learning over 1000 episodes as a measure of sample efficiency, averaged over 10 runs (F) Plot of flexibly modulated $\omega$ arbitration parameter over the learning duration averaged over 10 runs. This shows a transition from a higher Pavlovian bias to a more instrumental agent over episodes as learning about the environment reduces uncertainty (G) Comparison of different agents using a trade-off metric and to be used only for didactic purposes (using equation 16 and more details in Methods).

Appendix A.3 includes the performance comparison of agents with a suitable flexible $\omega$ and with fixed $\omega$ values on the three-route task. Appendix A.4 shows the results of a human experiment with subjects navigating a 3-route virtual reality maze similar to the simulated one.

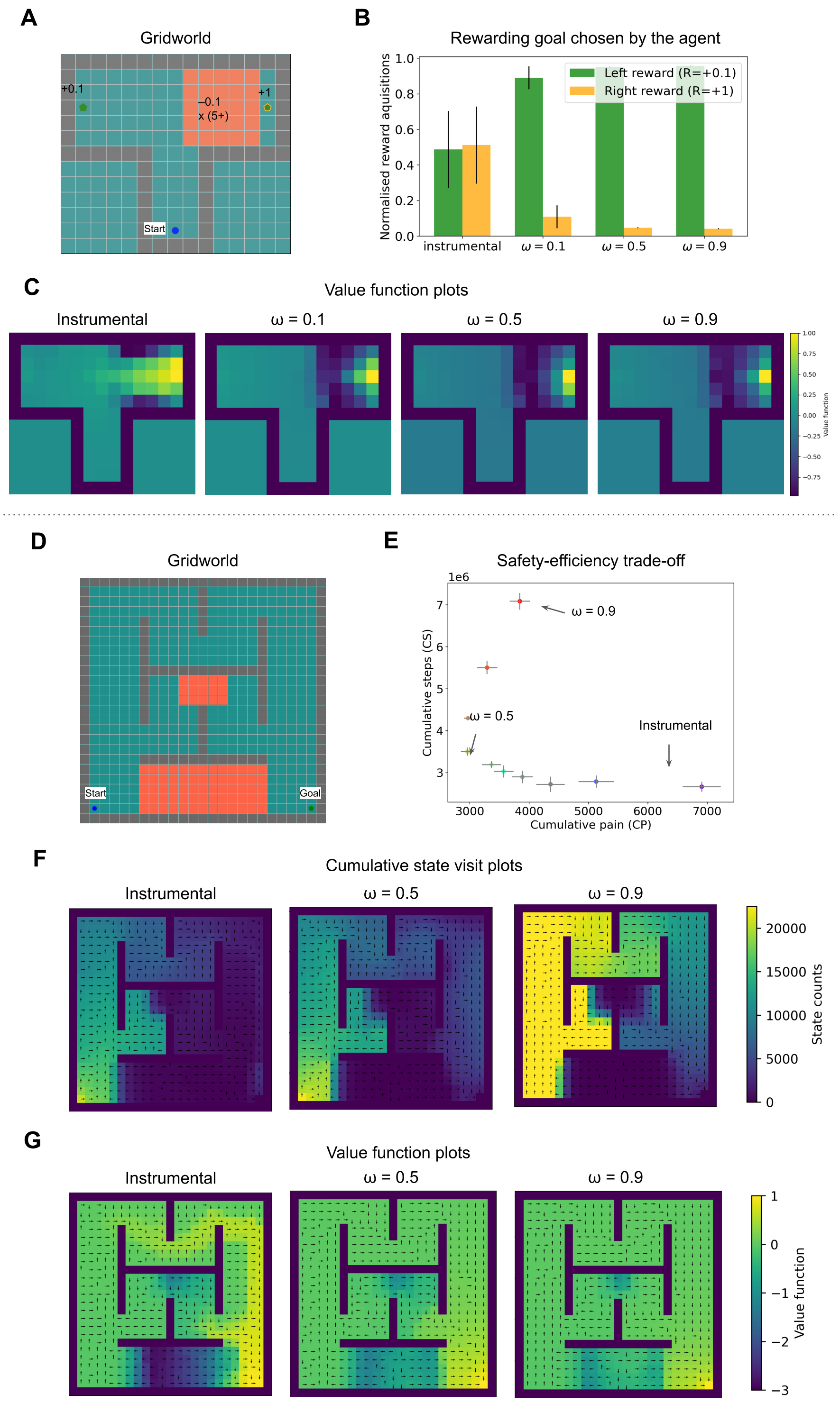


**Figure 3.** (A) T-maze grid world environment with annotated rewards and punishments. (B) Proportion of the rewarding goal chosen by the agent (C) Value function plots for $\omega = 0, 0.1, 0.5, 0.9$ shows diminished value propagation from the reward on the right (D) Grid world environment with three routes with varying pain (E) Cumulative steps required to reach the goal vs cumulative pain accrued by fixed $\omega$ agents ranging from $\omega = 0$ to $\omega = 0.9$ (F) State visit count plots for $\omega = 0, 0.5, 0.9$ i.e. instrumental and constant Pavlovian bias agents. (F)Value function plots for $\omega = 0, 0.5, 0.9$.

#### Experiment 3: Human approach-withdrawal conditioning is modulated by outcome uncertainty

Our first experiment showed the benefit of having a outcome uncertainty-based flexible $\omega$ arbitration scheme in balancing safety and efficiency, in a series of grid worlds. In this next experiment, we aimed to find basic evidence that humans employ such a flexible fear commisioning scheme. This is not intended as an exhaustive test of all predictions of the model, but to show in principle that there are situations in which a flexible, rather than fixed pavlovian influence, provides a good fit to real behavioural data. In line with our grid world simulations, we expected a Pavlovian bias in choices, but in addition to it, we also expected the a Pavlovian bias in reaction times.

Fig. 4 describes the trial protocol (Fig.4A), block protocol (Fig.4B) and experimental setup (Fig. 4C). We conducted a VR-based approach-avoidance task (28 healthy subjects, of which 14 females and average age 27.96 years) inspired by previous Go-No Go task studies for isolating Pavlovian bias, especially its contributions to misbehaviour [Guitart-Masip et al., 2012, Cavanagh et al., 2013, Mkrtchian et al., 2017a,b, Dorfman and Gershman, 2019, Gershman et al., 2021]. The subjects goal was to make a correct approach or withdrawal decision to avoid pain, with four different cues associated with different probabilities of neutral or painful outcomes. We expected the Pavlovian misbehaviour to cause incorrect withdrawal choices for cues where the correct response would be to approach. And in terms of reaction times, we expected the bias to slow down correct approach responses and speed up correct withdrawal responses. We explicitly attempted to change the outcome uncertainty or controllability, in a similar way to previous demonstrations Dorfman and Gershman [2019], but with controllability changing *within* the task. To do this, we set up two of the four cues to be uncontrollable in the first half (i.e. outcome is painful 50% of the times regardless of the choice), but which then become controllable in the second half (i.e. the correct choice will avoid the pain 80% of the times). We anticipated that the Pavlovian bias in choice and reaction times would be modulated along with the change in uncontrollability. The virtual reality environment improves ecological validity[Parsons, 2015] and introduces gamification, which is known to also improve reliability of studies[Sailer et al., 2017, Kucina et al., 2023, Zorowitz et al., 2023], which is important in attempts to uncover potentially subtle biases.

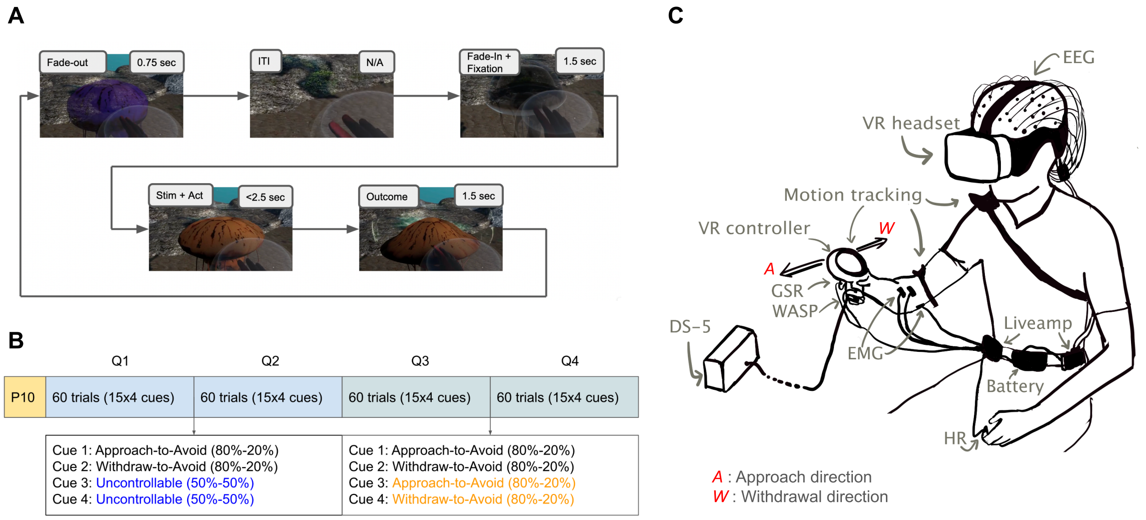


**Figure 4.** (A) Trial protocol: The participant is expected to take either an approach action (touch the jellyfish) or withdrawal action (withdraw the hand towards oneself) within the next 2.5 seconds once the jellyfish changes colour. The participant was requested to bring their hand at the centre of a bubble located halfway between the participant and the jellyfish to initiate the next trial where a new jellyfish would emerge. [Supplementary video] (B) Block protocol: First half of the trials had two uncontrollable cues and two controllable cues, and the second half had all controllable cues with aforementioned contingencies. The main experiment 240 trials were preceded by 10 practice trials which do not count towards the results. (C) Illustration of experimental setup VR: Virtual Reality, WASP: Surface electrode for electrodermal stimulation, DS-5: Constant current stimulator, GSR: galvanic skin response sensors, HR: Heart rate sensor, EMG: Electromyography sensors, EEG: Electroencephalogram electrodes, Liveamp: Wireless amplifier for mobile EEG.

Appendix A.5 includes behavioural results for the experiment data. Appendix A.6 includes group-level and subject-level (hierarchically fitted) model parameter distributions. Appendix A.7 includes tables of model parameters with LOOIC and WAIC values for all RL models and RLDDM models. We observed that all Rhat values were strictly less than 1.05 (most parameters were less than 1.01 and generally close to 1), indicating that the models converged.

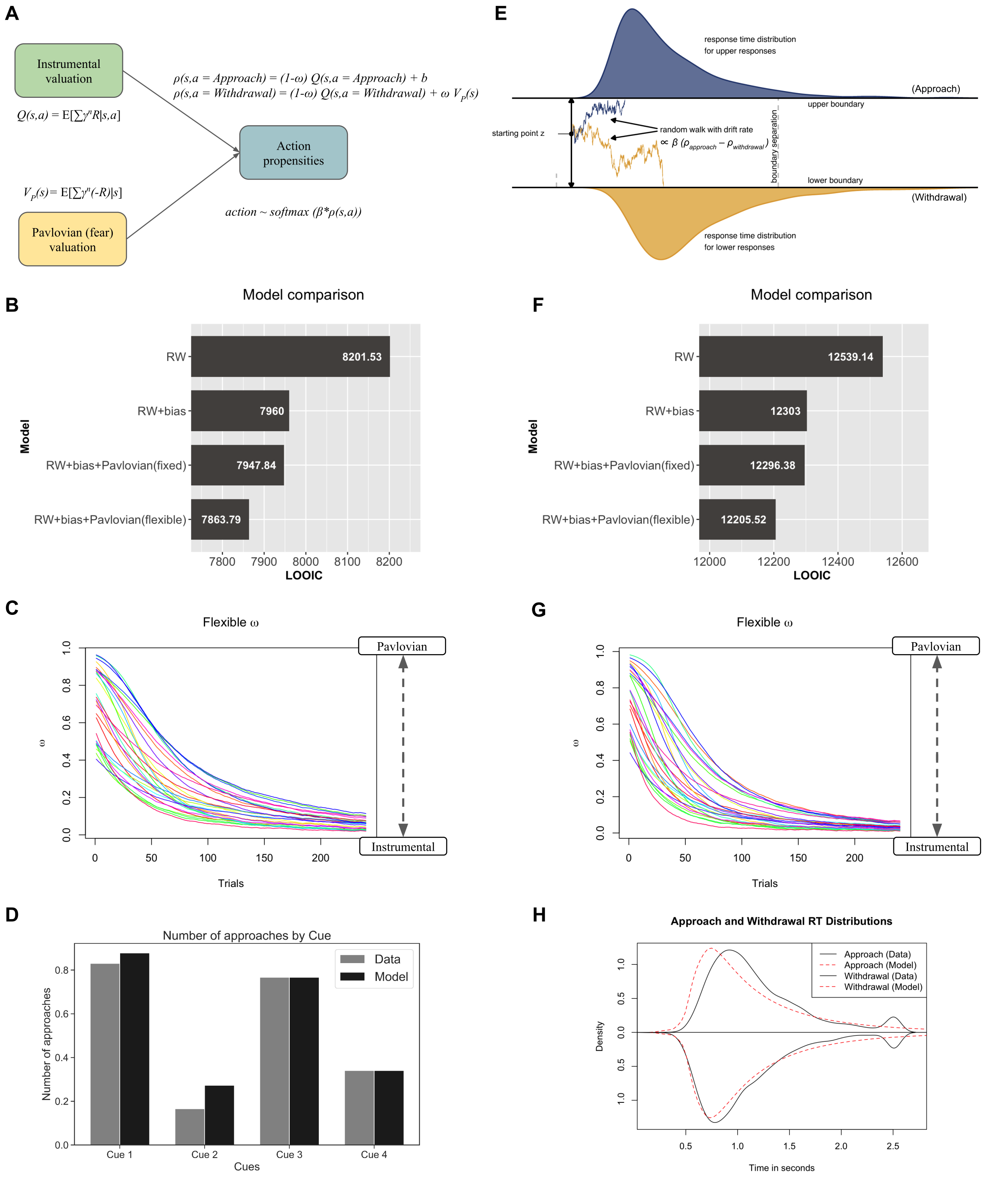


**Figure 5.** Left panels show choice model fit results using RL models. The right panels show choice and reaction times model fit results using RLDDM models. (A) Simplified RL model from Fig. 1 for the Approach-Withdrawal task (B) Model comparison shows that the model with flexible Pavlovian bias fits best to choices in terms of LOOIC (C) Flexible $\omega$ from the RL model over 240 trials for 28 participants. (D) Number of approaches aggregated over all subjects and all trials in data and model predictions by the RL model with flexible $\omega$. (normalized to 1) (E) Simplified illustration of RLDDM for the Approach-Withdrawal task, where the baseline bias $b$ and the Pavlovian bias $\omega V_p(s)$ is also included in the drift rate (The base figure is reproduced from Desch et al. [2022] with modifications) (F) Model comparison shows that the model with flexible Pavlovian bias fits best to choices and reaction times in terms of LOOIC (G) Flexible $\omega$ from the RLDDM over 240 trials for 28 participants (H) Distribution of approach and withdrawal reaction times (RT) aggregated over all subjects and trials in data and model predictions by the RLDDM with flexible $\omega$. The bump in RTs at 2.5 seconds is because of timeout (inactive approaches and withdrawals, please see Appendix A.5)

### MATERIALS AND METHODS

#### Instrumental learning and Pavlovian fear learning

We consider a standard reinforcement learning setting in an environment containing reward and punishments (pain). In each time step $t$, the agent observes a state ~~$s$~~ $s_t$ and selects an action ~~$a$~~ $a_t$ according to its stochastic policy ~~$\pi_t(s,a)$~~ $\pi_t(s_t,a_t)$ (i.e., the probability of selecting action $a_t = a$ in state $s_t = s$). The environment then makes a state transition from the current state ~~$s$~~ $s_t$ to the next state ~~$s'$~~ $s_{t+1}$ and the agent receives a scalar reward ~~$R \in (-\infty,+\infty)$.~~ $R_{t+1} \in (-\infty,+\infty)$. This represents that the scalar reward includes both positive rewards and negative rewards or punishments. We use the standard notation used by Sutton [2018].

In the Instrumental system, we define the value of taking action $a$ in state $s$ under a policy $\pi$, denoted as the action-value function $Q^{\pi}(s,a)$, as the expected return starting from $s$, taking the action $a$, and thereafter following policy $\pi$:

$$Q^{\pi}(s,a) = \mathbb{E}_{\pi}\left[\sum_{k=0}^{\infty}\gamma^{k}R_{t+k+1}|s_t = s, a_t = a\right] \tag{1}$$

And the

where $R_{t+k+1}$ is the scalar reward received $k$ time steps in the future, when evaluating the Q-values at timesteps $t$. The discount factor is $\gamma$ and the reward $k$ timesteps into the future is discounted by $\gamma^k$. The optimal action-value function is defined as $Q^*(s,a) = \max_{\pi} Q^{\pi}(s,a)$. Note this are purely instrumental Q-values and do not include the Pavlovian bias.

In addition to the instrumental system, we define a Pavlovian fear (i.e. punishment/pain) system over-and-above the instrumental system which makes it safer. The Pavlovian fear systems aims to increase the impetus of the pain-avoidance actions that minimize pain. For that we split the standard reward $R$ into pain (or punishment ) and only extract the punishment feedback signal $p \geq 0$:

$$p = -\min(R, 0), \tag{2}$$

We can similarly define a Pavlovian reward system trained on $max(R,0)$, however that's not relevant to the questions of this study so we will only focus on the arbitration between the instrumental (state-action based) model-free and Pavlovian (state-based) fear system. And we define the pain state-value $V_p(s)$ of the Pavlovian fear system as follows :

$$V_p^{\pi}(s) = \mathbb{E}_{\pi}\left[\sum_{k=0}^{\infty}\gamma^{k}p_{t+k+1}|s_t = s\right], \tag{3}$$

The subset of actions with the Pavlovian bias $A_p$ are arrived at using a pretraining in the same environemnt with only punishments and random starting points. $V_p(s)$ then bias this pretrained subset of actions $A_p$ according to equation 13.

Here onwards we will drop the time subscript for simplicity and write value update equations considering the state transtions from $s$ to $s'$. The Pavlovian fear state-value functions are updated as follows:

$$V_p(s) := V_p(s) + \alpha(p + \gamma V_p(s') - V_p(s)) \tag{4}$$

The instrumental value function for qualitative value plots is updated in an on-policy manner as follows (but is not used in the PAL algorithm):

$$V(s) := V(s) + \alpha(R + \gamma V(s') - V(s)) \tag{5}$$

And the Instrumental action-value functions are updated as follows:

$$Q(s,a) := Q(s,a) + \alpha(\delta) \tag{6}$$

where $\alpha$ is the learning rate and while using off-policy Q-learning (sarsamax) algorithm, the TD-errors are calculated as follows:

$$\delta = R + \gamma Q(s', \text{argmax}_{a'}(Q(s',a'))) - Q(s,a) \tag{7}$$

The equations above are valid for a general case and are used in grid-world simulations. For model-fitting purposes for the VR Approach-Withdrawal task, there is no next state $s'$, thus the equations reduce to a simpler form of the Rescorla-Wagner learning rule.

#### Action selection

Let $A$ be the action set. In the purely instrumental case, propensities $\rho(s,a)$ of actions $a \in A$ in state $s$ are the advantages of taking action $a$ in state $s$:

$$\rho(s,a) = Q(s,a) \tag{8}$$

And thus using softmax action selection with a Boltzmann distribution, the stochastic policy $\pi(a|s)$ (probability of taking action $a$ in state $s$) as follows:

$$\pi(a|s) = \frac{e^{(\rho(s,a)/\tau)}}{\sum_{a' \in A} e^{(\rho(s,a')/\tau)}} \tag{9}$$

where $\tau$ is the temperature that controls the trade-off between exploration and exploitation. For gridworld simulations, we use hyperbolic annealing of the temperature, where the temperature decreases after every episode $i$:

$$\tau(i) = \frac{\tau_0}{1 + \tau_k i} \tag{10}$$

Here $\tau_0$ is the initial temperature and $\tau_k$ controls the rate of annealing. This is to ensure the policy converges in large state spaces like a gridworld and follows previous studies [Elfwing and Seymour, 2017, Wang et al., 2018]. For model-fitting of the VR Approach-Withdrawal task we do not anneal it and keep it as a free parameter (inverse temperature $\beta = 1/\tau$) to be fitted to each participant, this is also consistent with previous literature [Guitart-Masip et al., 2012, Cavanagh et al., 2013, Dorfman and Gershman, 2019, Gershman et al., 2021] and several other works modelling Go-No Go tasks.

In case of a Pavlovian pain/fear system, let $A_p$ be the subset of actions in state $s$ which has the Pavlovian pain urges or impetus associated with it. These are usually a small set of species specific defensive reactions (SSDR). In the VR Approach-Withdrawal task we assume or rather propose it is the bias to withdraw from potentially harmful stimuli (in our case jellyfish). For the purpose of the gridworld simulations, these can either be hardcoded geographical controller moving away from harmful states or through a pretrained value-based controller [Dayan et al., 2006]. This work does not delve into the evolutionary acquisition of these biases, but one can derive the action subset $A_p$ from evolutionarily acquired value initializations which may also help avoid novel stimuli and is a direction for future work.

Thus after adding a Pavlovian fear system over and above the instrumental system, the propensities for actions are modified as follows:

$$\rho(s,a_n) = (1-\omega)Q(s,a_n); \text{ where } a_n \in A_n = A \setminus A_p. \tag{11}$$

$$\rho(s,a_p) = (1-\omega)Q(s,a_p) + \omega(V_p(s)); \text{ where } a_p \in A_p \tag{12}$$

The same can be compactly written as mentioned in the illustration (Fig. 1):

$$\rho(s,a) = (1-\omega)Q(s,a) + \omega(V_p(s,a)); \text{ where } V_p(s,a) = \mathbb{I}[a = a_p]V_p(s) \tag{13}$$

where $\omega$ is the parameter responsible for ~~Pavlovian-instrumental~~ Pavlovian-Instrumental transfer. These equations are constructed following the preceding framework by Dayan et al. [2006] which laid out the foundation for interplay between Pavlovian reward system and the instrumental system. $\mathbb{I}[\cdot] = 1 \forall a_p \in A_p$ and $\mathbb{I}[\cdot] = 0 \forall a_n \in A_n = A \setminus A_p$ following the succinct vectorised notation by [Dorfman and Gershman, 2019]. .

We have refered to this algorithm as the Pavlovian Avoidance Learning (PAL) algorithm in this paper. The equations above assume only Pavlovian fear system in addition to the an instrumental system, and the given equations would vary depending on if we add a Pavlovian reward system too. After this modification, the action selection probabilities are calculate in similar fashion as described in equation 9.

#### Uncertainty based modulation of $\omega$

We further modulate the parameter $\omega$ which is responsible for Pavlovian-instrumental transfer using perceived uncertainty in rewards. We use Pearce-Hall associability for this uncertainty estimation based on unsigned prediction errors [Krugel et al., 2009, Zhang et al., 2016, 2018]. We maintain a running average of absolute TD-errors $\delta$ (equation 7) at each state using the following update rule:

$$\Omega_{t+1} = (1 - \alpha_\Omega * \alpha)\Omega_t + \alpha_\Omega * \alpha \mid \delta \mid \tag{14}$$

where $\Omega$ is the absolute TD-error estimator, $\alpha$ is the learning rate for $Q(s,a)$ and $V(s)$ values as mentioned earlier and $\alpha_\Omega \in [0,1]$ is the scalar multiplier for the learning rate used for running average of TD-error. To obtain parameter $\omega \in [0,1]$ from this absolute TD-error estimator $\Omega \in [0,\infty)$, we scale it linearly using scalar $\kappa$ and clip it between $[0,1]$ as follows:

$$\omega_t = \min(\kappa\Omega_t, 1) \tag{15}$$

We note that the range values $\Omega$ takes largely depends on the underlying reward function in the environment and $\alpha_\Omega$. Thus we choose a suitable value of $\kappa$ for $\alpha_\Omega$ using gridsearch in each gridworld environment simulation to ensure that the Pavlovian system dominates in cases of high uncertainty and that the instrumental system starts to take control as uncertainty reduces. We aim to show that this flexible $\omega$ scheme is a viable candidate for arbitration between the two systems and addresses the safety-efficiency dilemma wherever it arises. The initial associability $\Omega_0$ is set to 0 in grid-world simulations as there is no principled way to set it. In the case of model-fitting for the VR Approach-Withdrawal task, $\Omega_0$, $\kappa$ and $\alpha_\Omega$ are set as free parameters fitted to each participant and instead of the TD-errors, we have the Rescorla-Wagner rule equivalent - punishment prediction errors (PPE) without any next state $s'$.

The action selection for RL models was performed using a softmax as per equation 9 with free parameter $\beta = 1/\tau$ and $\beta > 0$. The learning rule for RW models was:

$$Q(s,a) := Q(s,a) + \alpha(R - Q(s,a)) \tag{17}$$

where $R = -1$ in case of electric shocks or $R = 0$ in case of neutral outcome. Punishment $p$ can be defined as per equation 2 and thus $p = 1$ in case of electric shocks or $p = 0$ and the Pavlovian punishment value is calculated as per:

$$V_p(s) := V_p(s) + \alpha(p - V_p(s)) \tag{18}$$

$\alpha > 0$ is the learning rate and fitted as a free-parameter and note that here $V_p$ is always positive.

For RW+bias model,

$$\rho(s,a) = Q(s,a) + b; \text{ if } a = \text{Approach} \tag{19}$$

$$\rho(s,a) = Q(s,a); \text{ else} \tag{20}$$

Here $b \in (-\infty, +\infty)$ is the baseline bias, which if positive represents a baseline approach bias and if negative represents baseline negative bias and is not Pavlovian in nature.

For RW+bias+Pavlovian(fixed) and RW+bias+Pavlovian(flexible) models,

$$\rho(s,a) = (1-\omega)Q(s,a) + b; \text{ if } a = \text{Approach} \tag{21}$$

$$\rho(s,a) = (1-\omega)Q(s,a) + \omega(V_p(s)); \text{ if } a = \text{Withdrawal} \tag{22}$$

Here $\omega \in [0,1]$ is a free parameter for the RW+bias+Pavlovian(fixed) model. $\omega$ is not a free parameter for the RW+bias+Pavlovian(flexible) model, but computed as per equations 14 and 15 with free parameters $\Omega_0$ (initial associability), $\kappa$ (scaling factor for $\omega$) and $\alpha_\Omega$ (learning rate multiplier for associability).

### APPENDIX

#### (A.1) Robustness of the associability-based $\omega$ in gridworld simulations

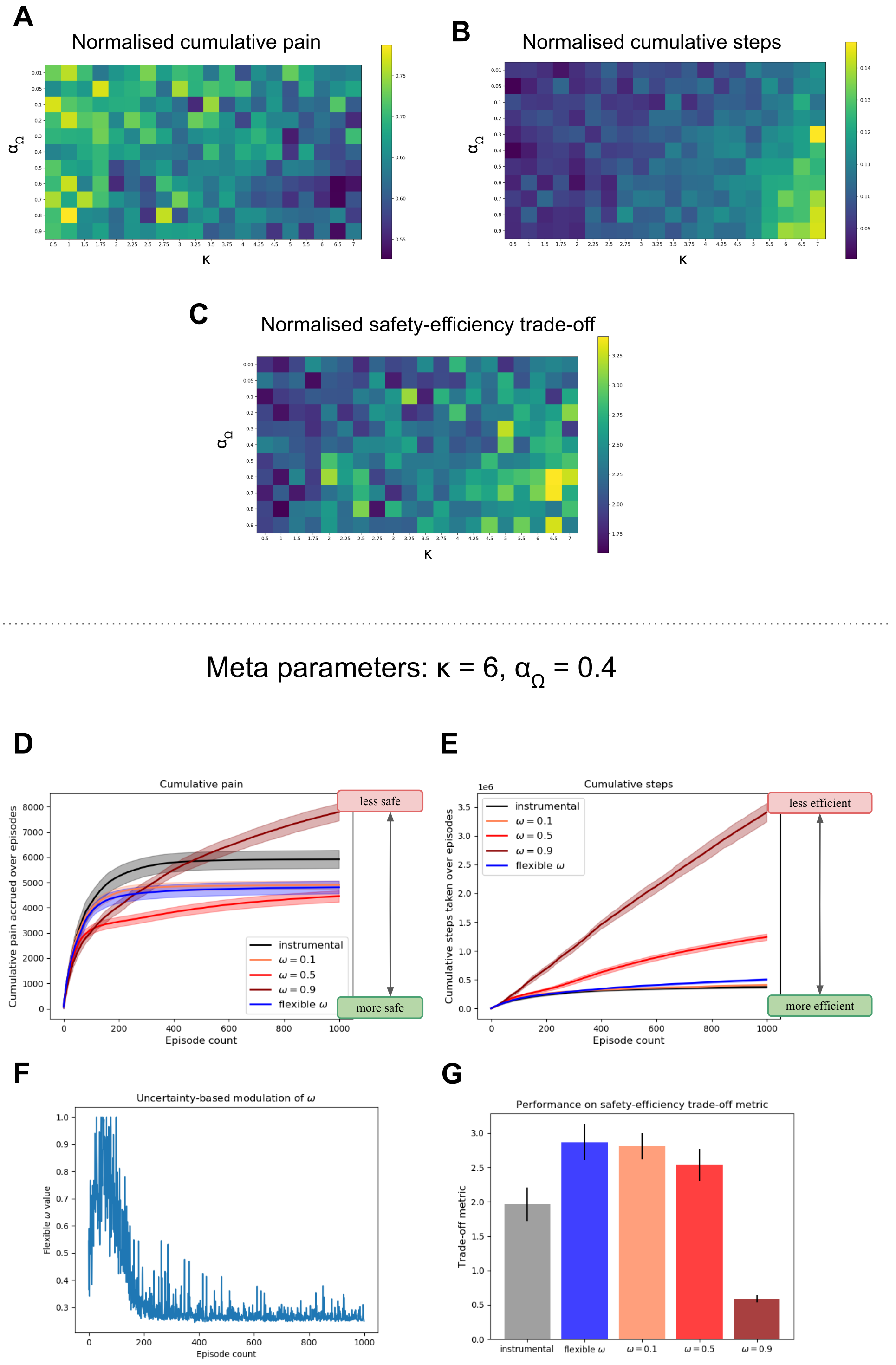


**Figure 6.** This figure shows the robustness of grid search for tuning the meta parameters for the associability-based $\omega$ in grid world simulations. We show that the results hold for a range of values close to the chosen meta-parameters. (A-C) Grid search results for the environment in Fig. 2 for varying $\kappa$ and $\alpha_{\Omega}$. (D-G) Results for another set of meta-parameters.

**(A.2) Flexible $\omega$ agent better adapts to reward relocation than a fixed $\omega$ agent.**

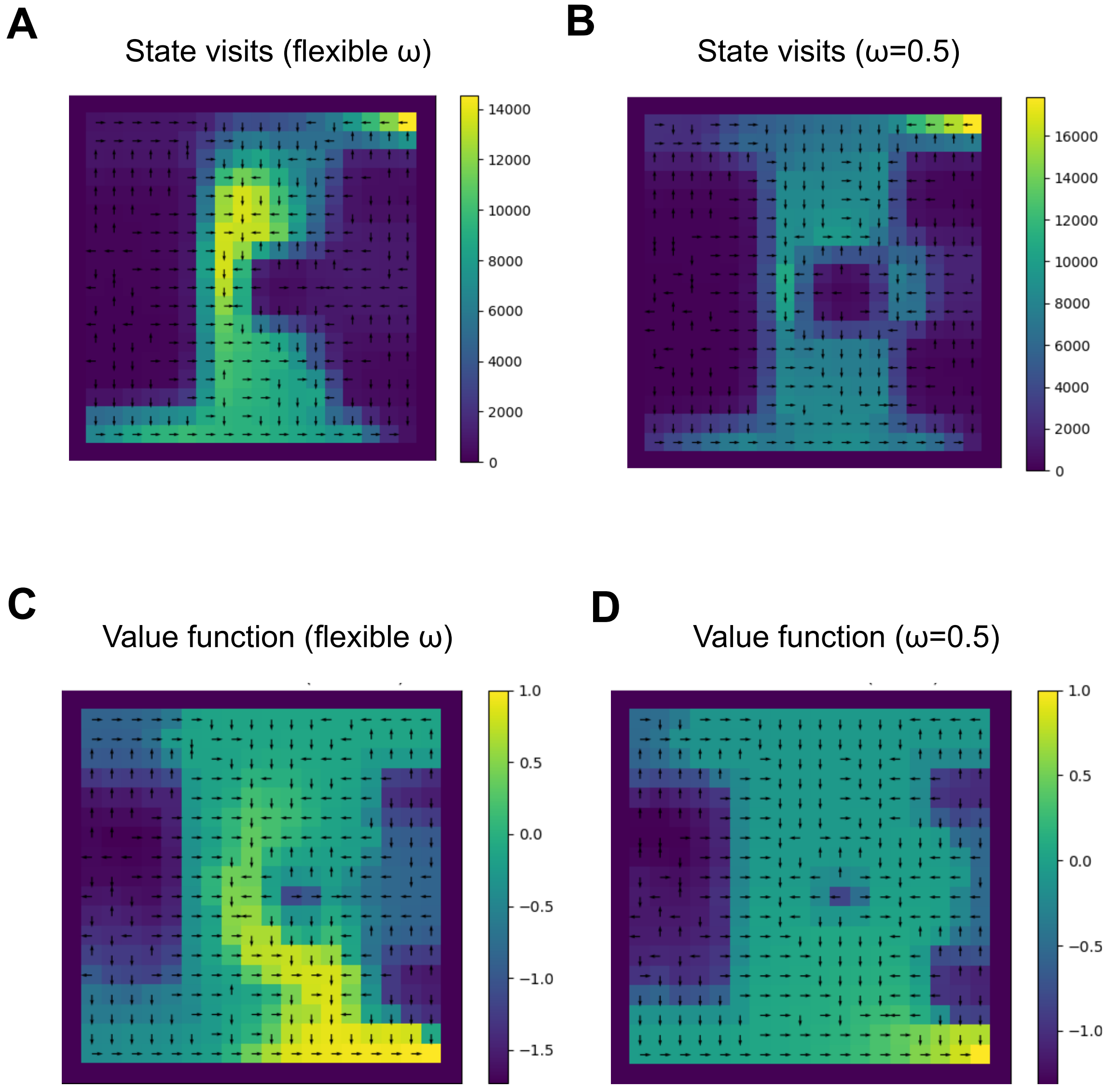


**Figure 7.** This figure shows cumulative state visit plots and value function plots of the flexible $\omega$ and fixed $\omega$ agents at the end of 1000 episodes when we relocate the reward goal from the bottom left corner (Fig. 2) to the bottom right corner on episode 500. Comparing state visit plots A & B and comparing value function plots C & D, we observe that persistent Pavlovian influence leads to persistent rigidity while the flexible fear commissioning scheme is able to efficiently locate the goal. We observe that unlike flexible $\omega$, constant $\omega = 0.5$ leads to diminished value propagation of the rewarding value (C & D).

##### (A.3) Solving the safety-efficiency trade-off in a range of grid world environments

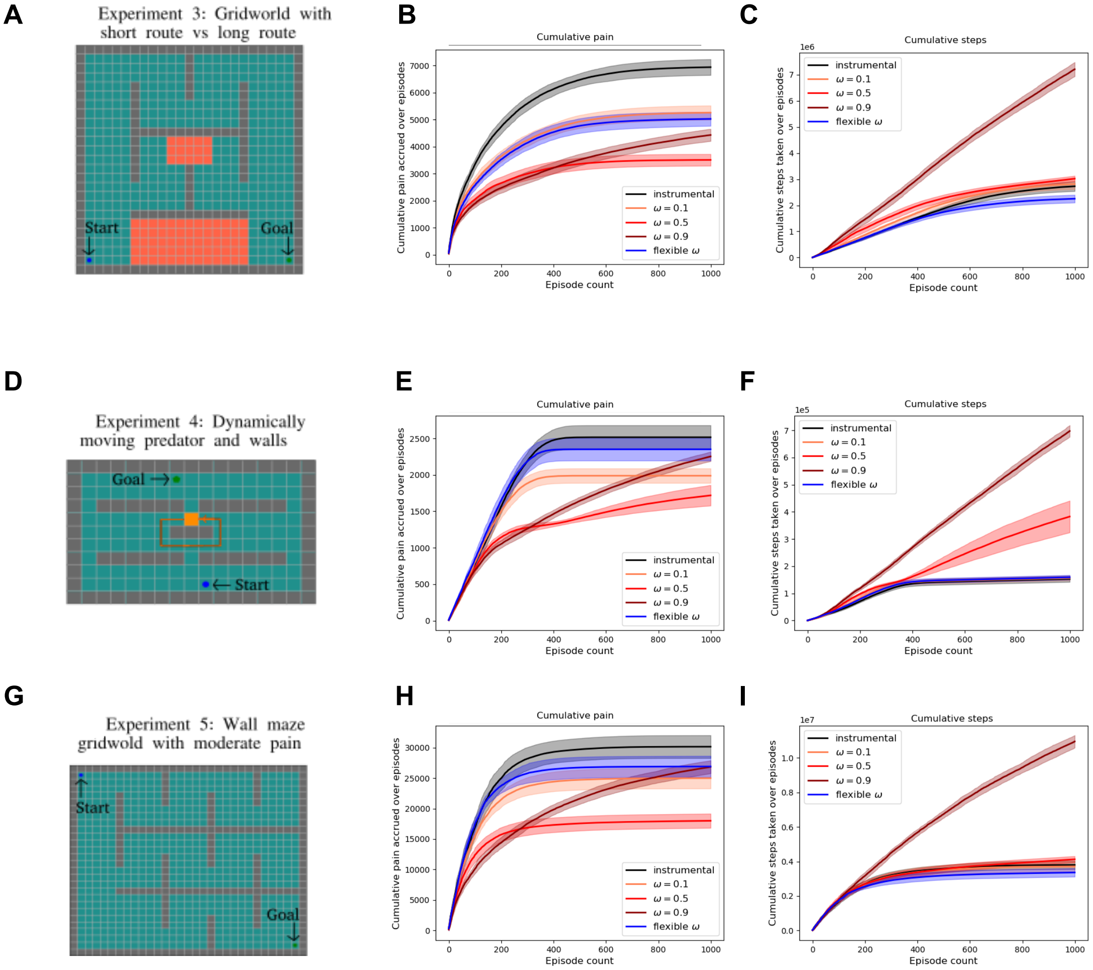


**Figure 8.** In this figure, we show the performance of fixed $\omega = 0.1, 0.5, 0.9$ and flexible $\omega$ agents on a range of grid world environments, namely (A) the three-route environment from Fig. 3, (D) an environment with a moving predator on routine path and (G) wall maze grid world from Elfwing and Seymour [2017]. Colliding with the predator results in a negative reward of -1 and catastrophic death (episode terminates). Otherwise, colliding with the walls results in moderate pain of 0.1, and the agent's state remains unchanged. The latter two are completely deterministic environments unlike the previous environments in the main paper. We show the safety-efficiency trade-off arises in these three environments as well and, there is a separate optimal fixed $\omega$ for each environment. Alternatively, there exists a flexible $\omega$ scheme for each environment that can solve the trade-off, suggesting that the brain may be calibrating $\omega$ flexibly.

#### (A.4) Human three-route virtual reality maze results

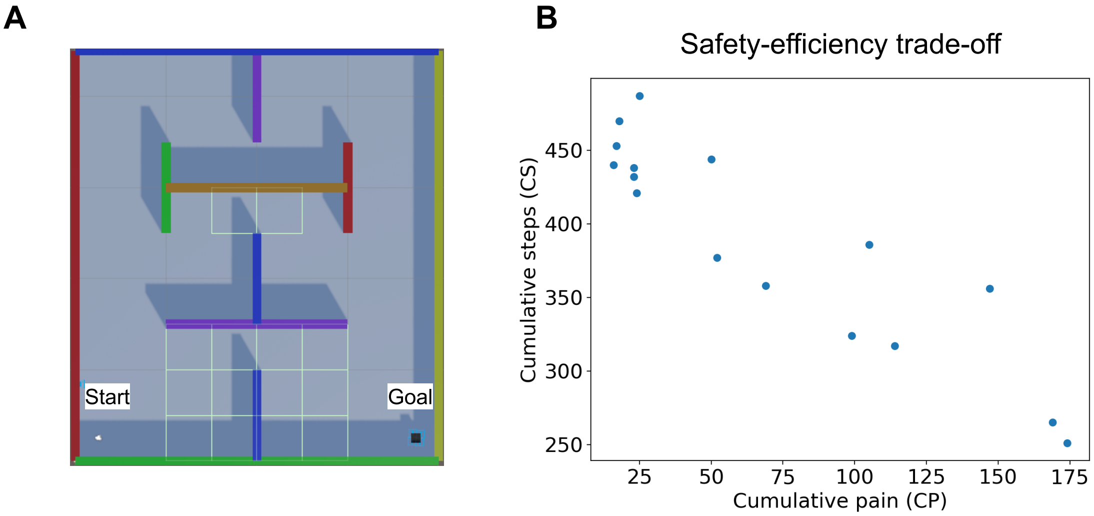


**Figure 9.** (A) Top-view of virtual reality (VR) maze with painful regions annotated by highlighted borders (B) Cumulative steps required to reach the goal vs cumulative pain acquired by participants over 20 episodes in the VR maze task. In this figure, we show the results of a VR maze replicating the three-route grid world environment from simulation results, however, it had fewer states and the participants were instructed to reach the goal which was visible to them as a black cube with "GOAL" written on it. In order to move inside the maze participants had to physically rotate in the direction they wanted to move and then press a button on the joystick to move forward in the virtual space. Thus the participant did not actually walk in the physical space but did rotate up to 360 degrees in physical space. The painful regions were not known to the participants but they were aware that some regions of the maze may give them painful shocks with some unknown probability. Walking over the painful states in the VR maze, demarcated by grid borders (see A) in the potentially shocked them with 75% probability while ensuring 2 seconds of pain-free interval between two consecutive shocks. Participants were not given shocks with 100% probability as that would be too painful for participants due to the temporal summation effects of pain. The participants engaged in 20 episodes of trials and were aware of this before starting the task and were free to withdraw from the experiment at any point. 16 participants (11 female, average age 30.25 years) were recruited and were compensated adequately for their time. The pain tolerance was acquired similarly to the Approach-Withdrawal task. (B) All participant trajectories inside the maze were discretized into an 8x9 (horizontal x vertical) grid. Entering a 1x1 grid section counted incremented the cumulative steps (CS) count. Upon receiving the shocks, the cumulative pain (CP) count was incremented. CP and CS over 20 episodes were plotted against each other to observe the trade-off. A limitation of this experiment is that it reflects the constraints of the grid world and future experiments are necessary to show the trade-off in a range of environments.

#### (A.5) Behavioural results from Approach-Withdrawal VR task

We observe a baseline approach bias through significant asymmetry in average number of cumulative approaches and withdrawals (Fig. 10A) and we consider the few inactive approaches and inactive withdrawals due to timeout as approaches and withdrawals respectively (Fig. 10B).

We consider a couple of model-free metrics of Pavlovian withdrawal bias prior to model fitting. The withdrawal bias metric on choices for two cues (say, cues X and cue Y) calculated as follows:

$$\text{Choice bias metric(cue X, cue Y)} = \text{\% withdrawal choices on `cue X'} - \text{\% approach choices on `cue Y'} \quad (24)$$

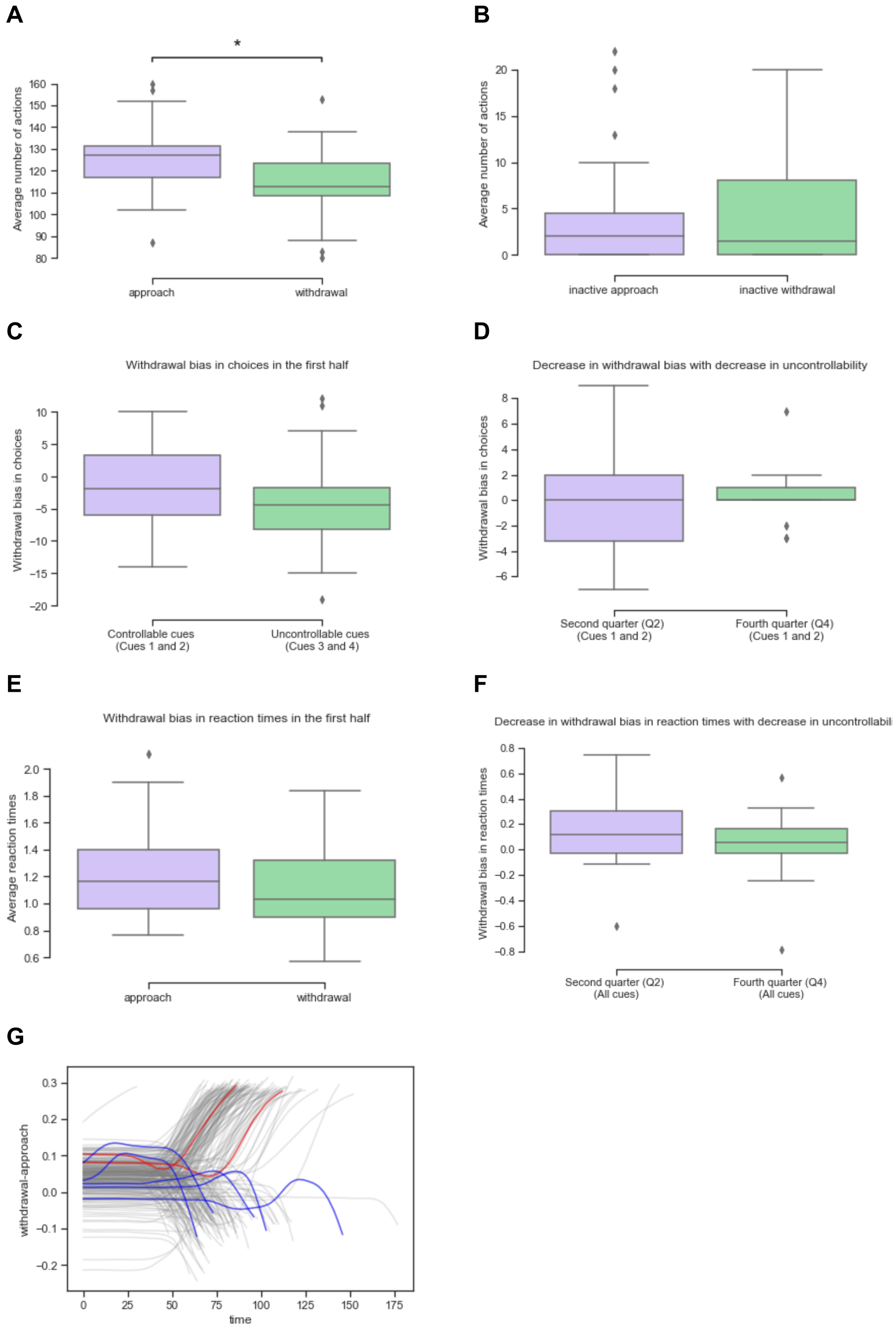


**Figure 10.** (A) Asymmetry in average approach and withdrawal responses over all subjects showing a baseline approach bias - MannwhitneyuResult(statistic=597.0, pvalue=0.0003981)(B) Incomplete approaches and incomplete withdrawals that were counted as approaches and withdrawals respectively. (C) Withdrawal bias in choices in the first half with uncontrollable cues - MannwhitneyuResult(statistic=492.5, pvalue=0.05037) (D) Decrease in withdrawal bias in choice with decrease in uncontrollability - MannwhitneyuResult(statistic=350.5, pvalue=0.76108) (E) Withdrawal bias in reaction times in the first half with uncontrollable cues - MannwhitneyuResult(statistic=475.0, pvalue=0.08820) (F) Decrease in withdrawal bias in reaction times with decrease in uncontrollability - MannwhitneyuResult(statistic=456.0, pvalue=0.14903) (G) Change-of-mind trials observed in motor data.

#### (A.6) Group and subject level parameter distributions of RL and RLDDM models

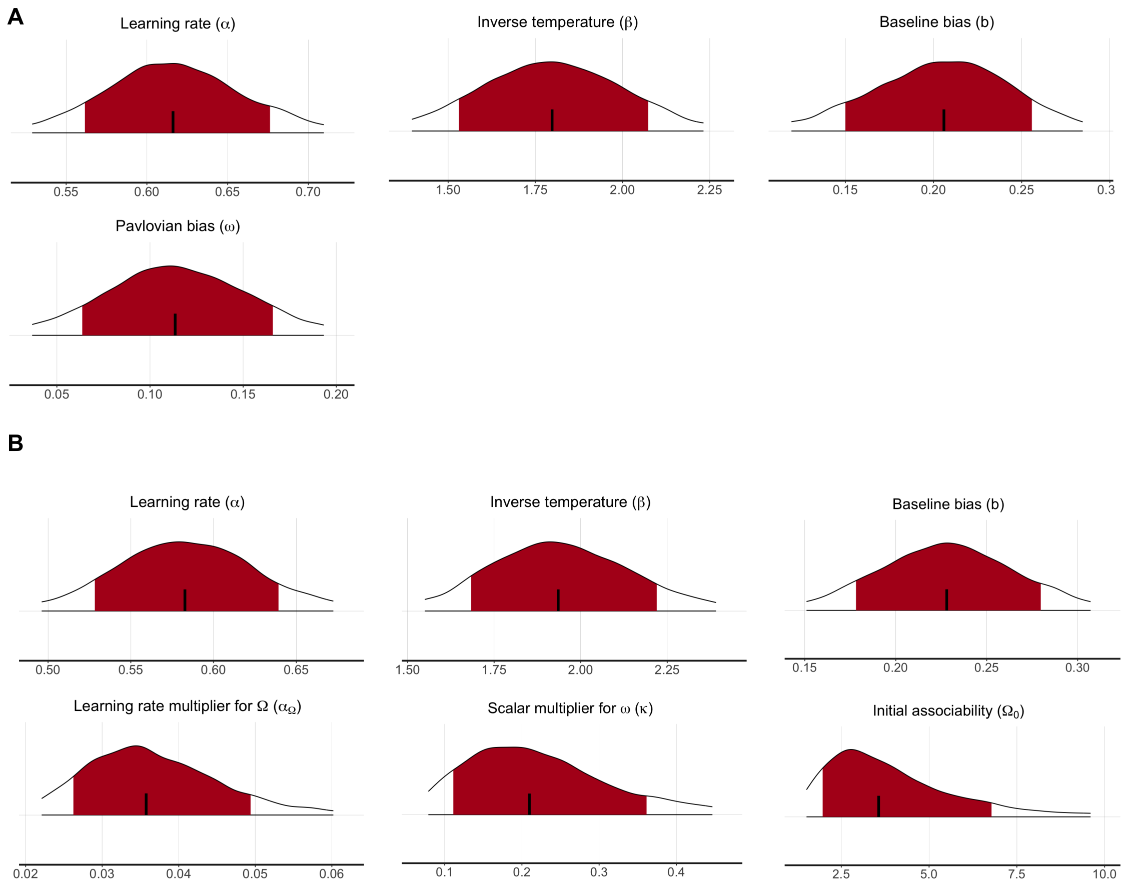


**Figure 11.** Group-level parameter distributions from (A) the RL model (M3) with fixed $\omega$ and (B) the RL model (M4) with flexible $\omega$. Shaded red regions denote 95% confidence intervals.

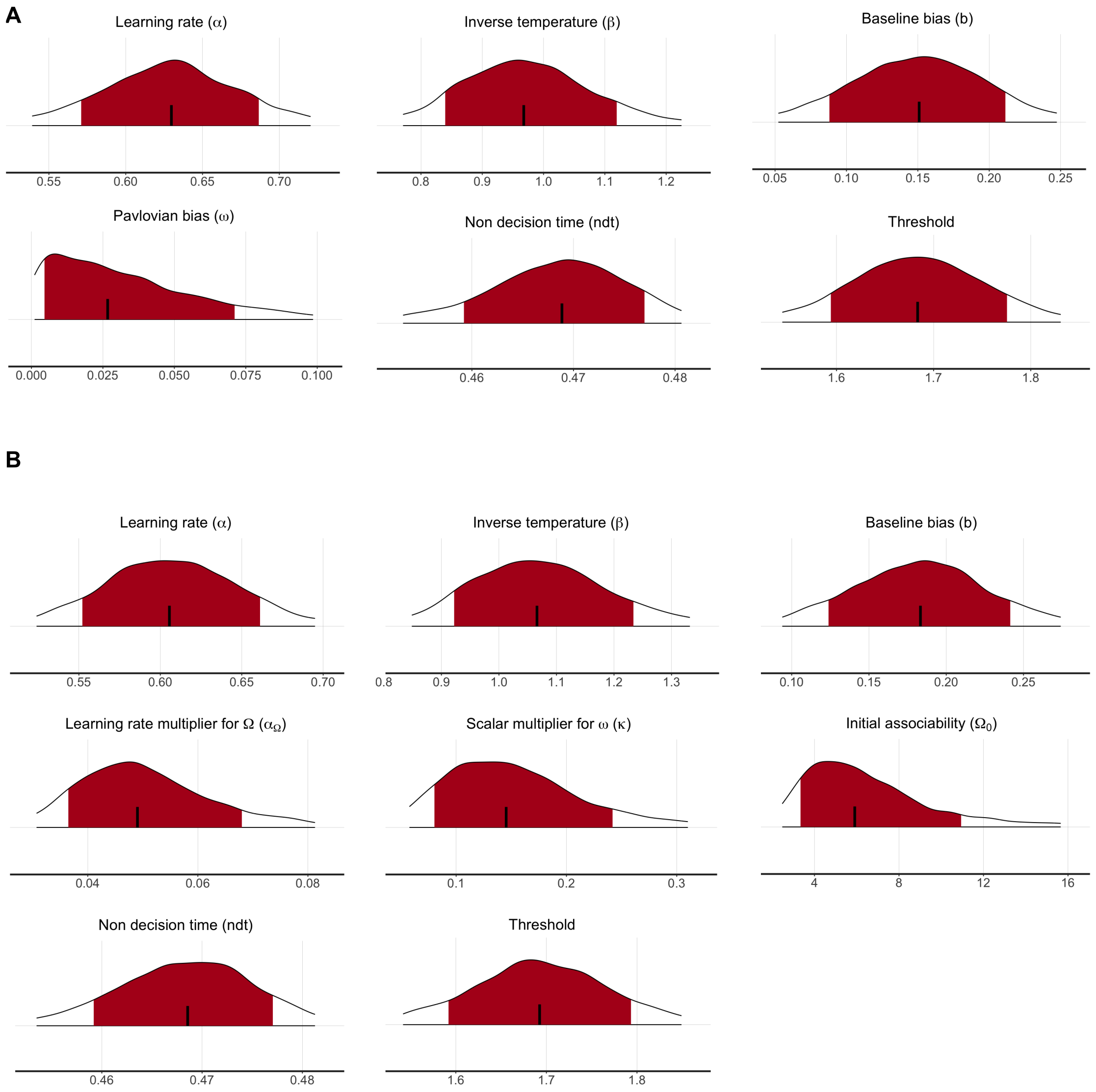


**Figure 12.** Group-level parameter distributions from (A) the RLDDM model (M3) with fixed $\omega$ and (B) the RLDDM model (M4) with flexible $\omega$. Shaded red regions denote 95% confidence intervals.

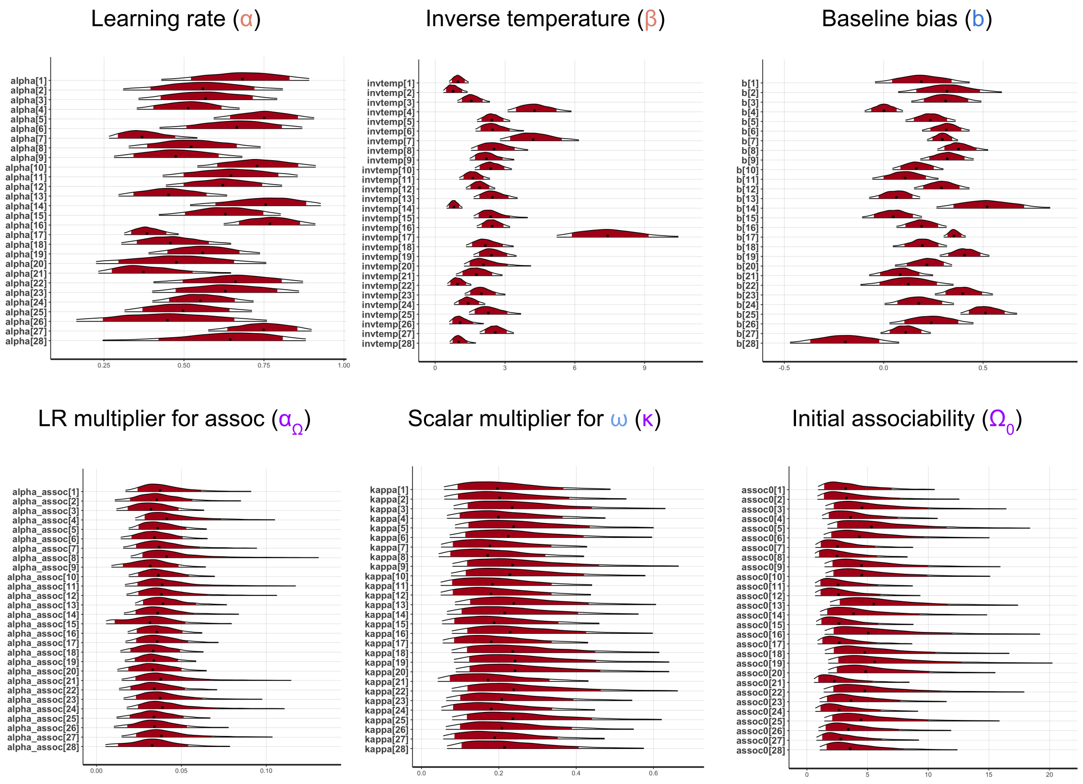


**Figure 13.** Subject-level parameter distributions from (A) the RL model (M3) with fixed $\omega$ and (B) the RL model (M4) with flexible $\omega$. Shaded red regions denote 95% confidence intervals.

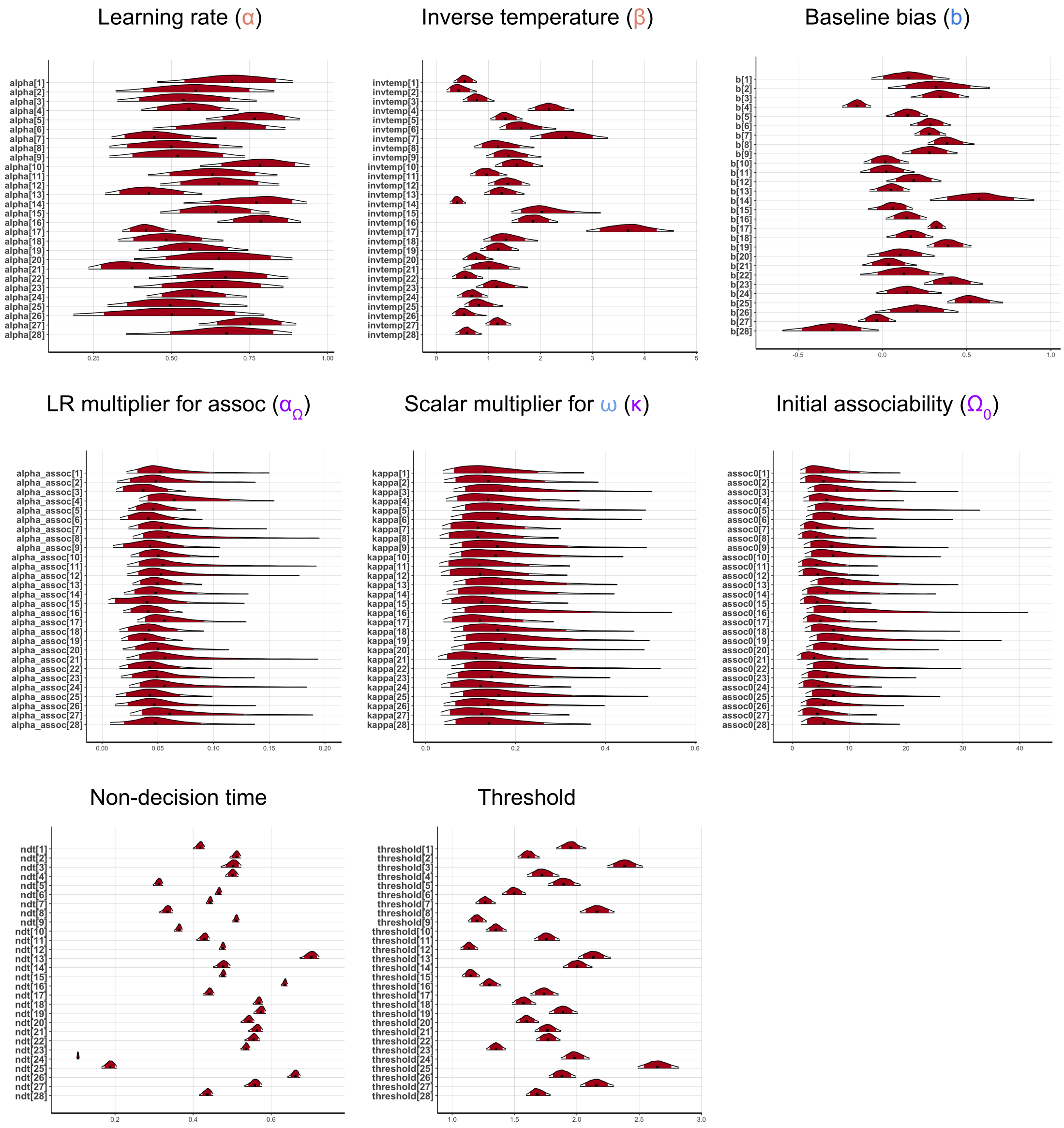


**Figure 14.** Subject-level parameter distributions from (A) the RLDDM model (M3) with fixed $\omega$ and (B) the RLDDM model (M4) with flexible $\omega$. Shaded red regions denote 95% confidence intervals.

#### (A.7) RL and RLDDM model parameters and model comparison tables

Please refer to Table 1 and Table 2.

**Table 1.** Model comparison results for RL models

| Model | Free Parameters | LOOIC | WAIC |
|---|---|---|---|
| M1 | $\alpha,\beta$ | 8201.53 | 8182.15 |
| M2 | $\alpha,\beta,b$ | 7960.00 | 7926.73 |
| M3 | $\alpha,\beta,b,\omega$ | 7947.84 | 7918.20 |
| M4 | $\alpha,\beta,b,\alpha_\Omega,\kappa,\Omega_0$ | 7863.79 | 7830.18 |

**Table 2.** Model comparison results for RLDDM model

| Model | Free Parameters | LOOIC | WAIC |
|---|---|---|---|
| M1 | ndt, threshold, $\alpha,\beta$ | 12539.14 | 12495.44 |
| M2 | ndt, threshold, $\alpha,\beta,b$ | 12303.00 | 12247.63 |
| M3 | ndt, threshold, $\alpha,\beta,b,\omega$ | 12296.38 | 12247.22 |
| M4 | ndt, threshold, $\alpha,\beta,b,\alpha_\Omega,\kappa,\Omega_0$ | 12205.52 | 12164.14 |

#### (A.8) Model predictions: Adapting fear responses in a chronic pain gridworld

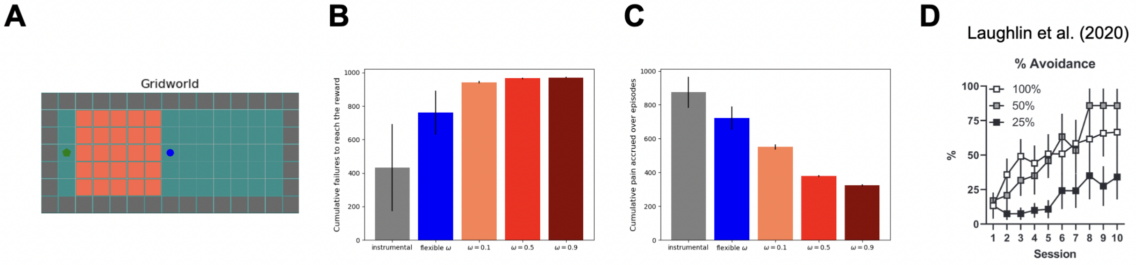


**Figure 15.** Pavlovian-instrumental interactions are invoked in a popular model of chronic pain, in which excessive Pavlovian fear of movement is self-punitive in a context in which active avoidance would reduce pain [Meulders et al., 2011, Crombez et al., 2012]. (A) Grid world with a start at the centre (blue) and goal at the left end (green, operationalises this. We augment the action set to include an additional "immobilize" action, to the action set, resulting in no state change and repeated rewards. An upper bound of 100 steps per episode is set; exceeding it leads to a painless death and episode restart. (B) Cumulative failures to reach the goal as a measure of efficiency. With a constant Pavlovian fear influence, the agent struggles to complete episodes, resembling effects seen in rodent models of anxiety [Laughlin et al., 2020] (C) Cumulative pain accrued as a measure of safety. In clinical terms, the agent remains stuck in a painful state, contrasting with an instrumental system that can seek and consume rewards despite pain. Flexible parameter $\omega$ ($\kappa = 3, \alpha_\Omega = 0.01$) allows the agent to overcome fear and complete episodes efficiently, demonstrating a safety-efficiency dilemma. The flexible $\omega$ policy outperforms fixed variants, emphasising the benefits of adapting fear responses for task completion. (D) Results from Laughlin et al. [2020] showing 25% of the (anxious) rats fail signalled active avoidance task due to freezing. GIFs for different configurations: pure instrumental agent, adaptively safe agent (flexible $\omega$) and maladaptively safe agent (constant $\omega$) can be found here.

#### (A.9) Neurobiology of Pavlovian contributions to bias avoidance behaviour

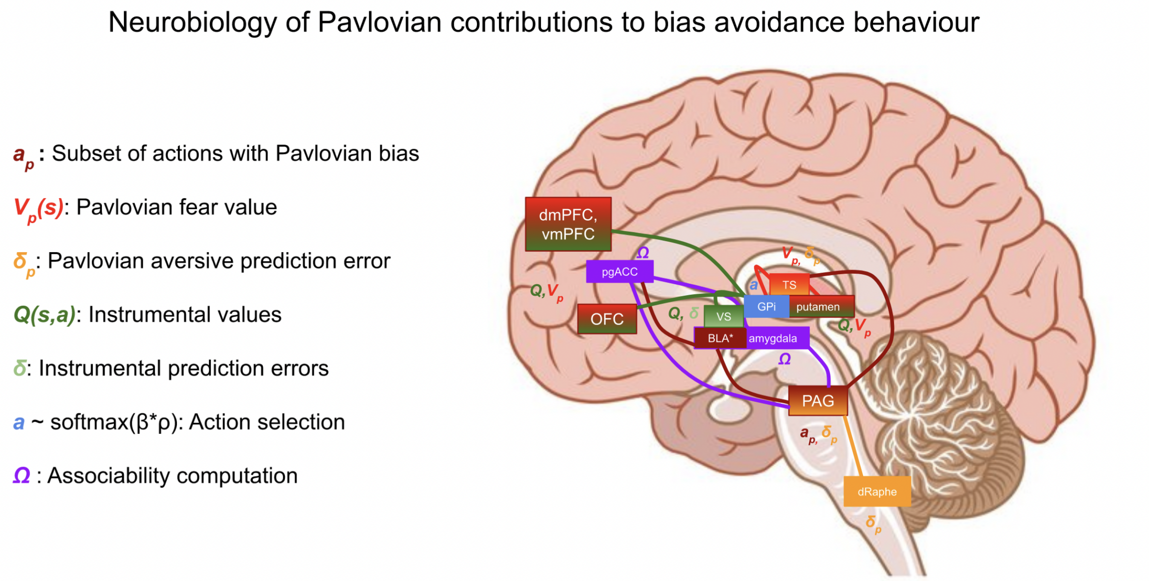


**Figure 16.** An overview of neurobiological substrates for the proposed PAL model based on relevant prior literature.
